# First Complete Genome Sequence of the Novel Lineage G-IX (BD-18) of FMDV Serotype Asia1

**DOI:** 10.1101/776518

**Authors:** A. S. M. Rubayet Ul Alam, M. Rahmat Ali, Md. Al Amin, Mohammad Anwar Siddique, Munawar Sultana, M. Anwar Hossain

## Abstract

Foot-and-Mouth Disease (FMD) has been a major threat to livestock worldwide, and is caused by FMD Virus (FMDV) existing as seven serotypes (A, O, C, Asia1 and SAT 1-3), each having variability into topotypes, genetic lineages, sublineages, and strains. Three serotypes are circulating in Indian subcontinent with the widespread distribution of serotype O, whereas serotype A and Asia1 are restricted to certain geographical regions. During 2017-2018, the Sindh-08 lineages of Asia1 serotype was reported from Pakistan, however, in 2018, a novel circulatory foot-and-mouth disease virus serotype Asia1 BD-18 (G-IX) lineage containing a unique mutation has emerged in Bangladesh. The first complete genome of the Asia1/ASIA/G-IX novel lineage strain, isolated from Bangladesh, is reported here. Amino acid substitutions at critical antigenic sites of capsid were identified compared to genome of existing vaccine strain (IND/63/72), and contemporary FMDV serotype A isolate of Bangladesh.

## INTRODUCTION

Foot-and-Mouth Disease (FMD), a highly infectious disease of livestock, causes global economic loss through impeding animal productivity and trade between countries. FMD Virus (FMDV), the etiological agent, is an RNA virus of *Aphthovirus* genus under *Picornaviridae* family with high mutation rate, hence generating various topotypes, lineages/groups and sublineages within its seven serotypes O, A, C, Asia1 and SAT (1-3) (Brito et al., 2017). In Bangladesh, FMD outbreaks due to three FMDV serotypes O, A and Asia1 were reported regularly since 2009, and two novel sublineages, namely Ind2001BD1 and Ind2001BD2, within O/ME-SA/Ind2001 lineage emerged in recent years (Siddique et al., 2018; Nandi et al., 2015;). Previously, we reported the complete genome sequences of all three circulatory serotypes, and three representative isolates of Ind2001d (BAN/NA/Ha-156/2013), Ind2001BD1 (BAN/GO/Ka-236(Pig)/2015), and BAN/BO/Na-161/2013 in Bangladesh (Sultana et al., 2014, Ullah et al., 2015, Ali et al., 2017, Alam et al., 2019). Recently, a novel Asia1 strain emerged in Bangladesh in 2018 (Ali et al. 2019) which was under lineage G-IX and differs substantially in VP1 coding region. We endeavor here to characterize its complete genome for further clarification at the genomic scale and finding the distinguishing features of the strain considering complete genomes of other lineages and related Bangladesh strain.

## MATERIALS AND METHODS

### Sample Collection and Processing

We collected the tongue epithelium sample of an infected cow at BGB farm, Pilkhana, Dhaka, Bangladesh on 24 January 2018. The sample was collected by registered veterinary doctor with the consent of the authority, and the Animal Experimentation Ethical Review Committee (AEERC) under Faculty of Biological Sciences, University of Dhaka approved the protocol for sample collection. The samples were immediately transported to the laboratory maintaining cold chain after collection and stored at −80°C before RNA extraction. The virus was isolated in BHK-21 cell line and viral RNA was extracted from infected cell culture supernatant of passage 3.

### RNA extraction and cDNA synthesis

RNA was extracted using Maxwell® 16 LEV RNA Cartridge in automated Maxwell® 16 nucleic acid extraction instrument following manufacturer’s protocol (Promega, USA). The extracted RNA was stored at −80°C until further use. The extracted RNA was reverse transcribed into cDNA using GoScript™ Reverse Transcription System (Promega, USA) as per the manufacture’s instruction. Briefly, primer/ RNA mix was prepared by mixing 5 µl of extracted RNA with 2 μl of Random primer, 2 μl of Oligo(dT)_15_ primer and 1 μl of Nuclease-free water (total volume 10 μl). Then the mixture was heated at 70°C in heat block for 5 minutes. Afterwards, the mixture was quickly chilled on ice for 5 minutes. The reverse transcription reaction mix was prepared by combining the components of the GoScript™ Reverse Transcription System in a sterile 1.5ml micro centrifuge tube keeping on ice. Forty (40) μl of reaction mix was prepared for each cDNA synthesis reaction to be performed.

### Genome amplification with overlapping primer sets

The entire genome was amplified by PCR using GoTaq 2× Hot Start Colorless Master Mix (Promega, USA) using overlapping primer sets (Ali et al., 2017). Afterwards, we purified PCR amplicons using the Wizard® SV Gel and PCR Clean-Up System (Promega, USA) and sequenced purified PCR products with BigDye® Terminator v3.1 cycle sequencing kit (Applied Biosystems®, USA) according to manufacturer’s instructions followed by analysis of data in ABI Genetic Analyzer (Applied Biosystems®, USA). Afterwards, the raw sequence data were assembled using SeqMan version 7.0 (DNASTAR, Inc., Madison, WI, USA), and we compared assembled sequences with other entries from NCBI GenBank using BLAST search to reveal the serotype level identification of the samples before submitting into NCBI GenBank.

### Phylogenetic and Diversity Analysis

The CDS from all the available FMDV Asia1 serotype of the NCBI database were taken into the phylogenetics analysis. Both heuristic program in MEGA7 (Kumar, Stecher, & Tamura, 2016) and jModelTest (version 2.1.10) package were used for model selection by computing likelihood score out of 88 models and best fitted model was selected using lowest Akaike Information Criterion (AIC) and Bayesian Information Criterion (BIC) values, which pointed out GTR (generalised time reversible) with a discrete gamma distribution of 1.46 with 4 categories and invariant proportion of 0.42 as the best fitted model to the aligned data set. We reconstructed CDS based phylogeny by maximum likelihood method using the assorted dataset in MEGA7 under on general time-reversible model of substitution with gamma distributed rate variation among sites with 1,000 pseudo-replicates for checking the robustness of the tree topology.

### Mutational Analysis

We checked the amino acid substitutions in the VP (1-3) region for our isolate against the present vaccine strain of serotype Asia1. Moreover, the protein variability of the proteins are checked against the BAN/TA/Ma-167/2013 (MF782478) strain, which is the closely related virus isolate in BLASTp analysis and belong to G-VIII.

## RESULTS AND DISCUSSION

The complete genome sequence of FMDV Asia1/ASIA/G-IX (BD-18) lineage isolate has been deposited in the NCBI GenBank database under the accession no. MN366244. The complete genome is 8,197 nucleotides(nt) in length with 54% GC-content including a 1089-nt lengthy 5’-untranslated region (UTR) with 18-nt poly(C) tract, and a 3’-UTR of 118-nt trailed by ≥24-nt poly(A) tail. The 6,990-nt long open reading frame (ORF) encodes 2,330 amino acids. The CDS based phylogeny corroborated with the VP1 phylogeny, which clustered the virus into distinct clade from other established lineages (G-I to G-VIII) (Figure 1). The phylogenetic tree showed that the strain is most closely related to another strain of Bangladesh, BAN/TA/Ma-167/2013 of G-VIII that is previously circulating in Bangladesh.

**Figure 1.**
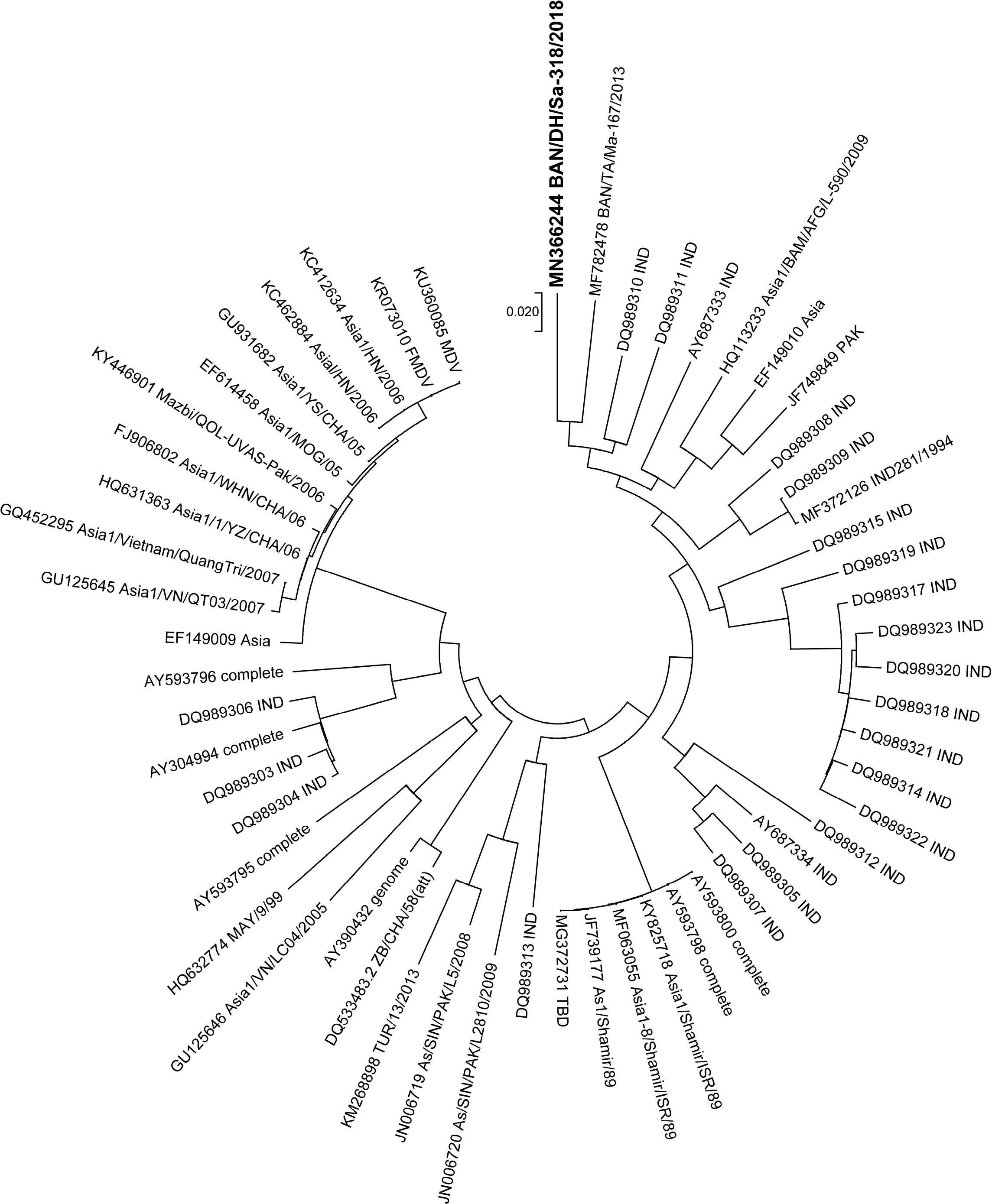
Phylogenetic tree representing the position of BAN/DH/Sa-318/2018 strain with other complete genomes of Asia1. The ML tree showed that the newly emerged strain clustered with the other previously circulated strain of Asia1 in Bangladesh, BAN/TA/Ma-167/2013.

The maximum identity of complete genomic nucleotide and CDS encoded amino acid sequence of BAN/DH/Sa-318/2018 was determined at 92.5% and 98%, respectively, with other virus strains of serotype Asia1. Compared to the vaccine strain (O/India/63/72, GenBank accession no. AY304994), BAN/DH/Sa-318/2018 capsid proteins (VP1-VP3) has thirteen crucial mutations at three five antigenic sites (loop regions), respectively (Table 1). This virus strain might pose a threat to vaccine escape since it has been changed substantially in capsid region.

**Table 1.**
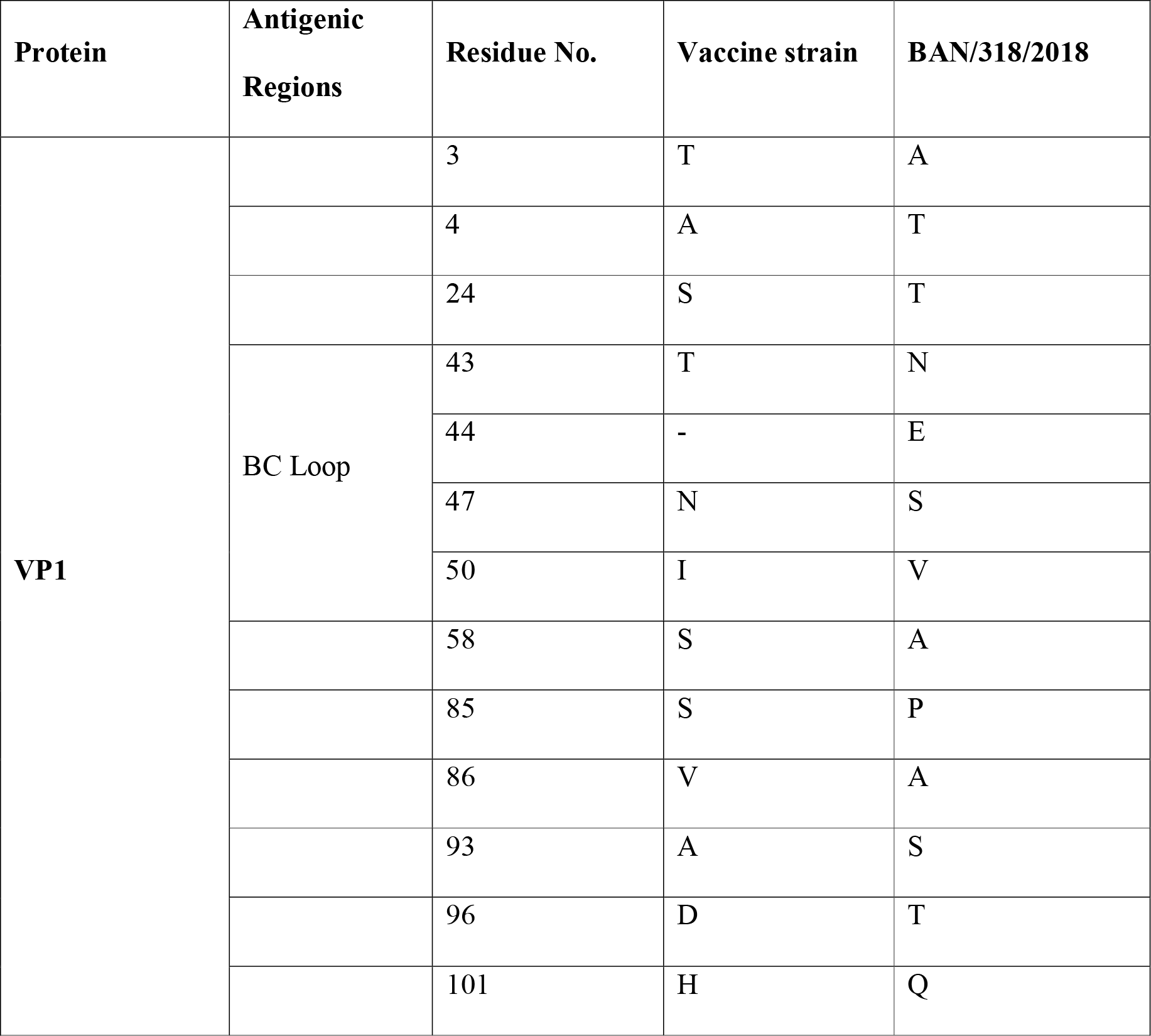
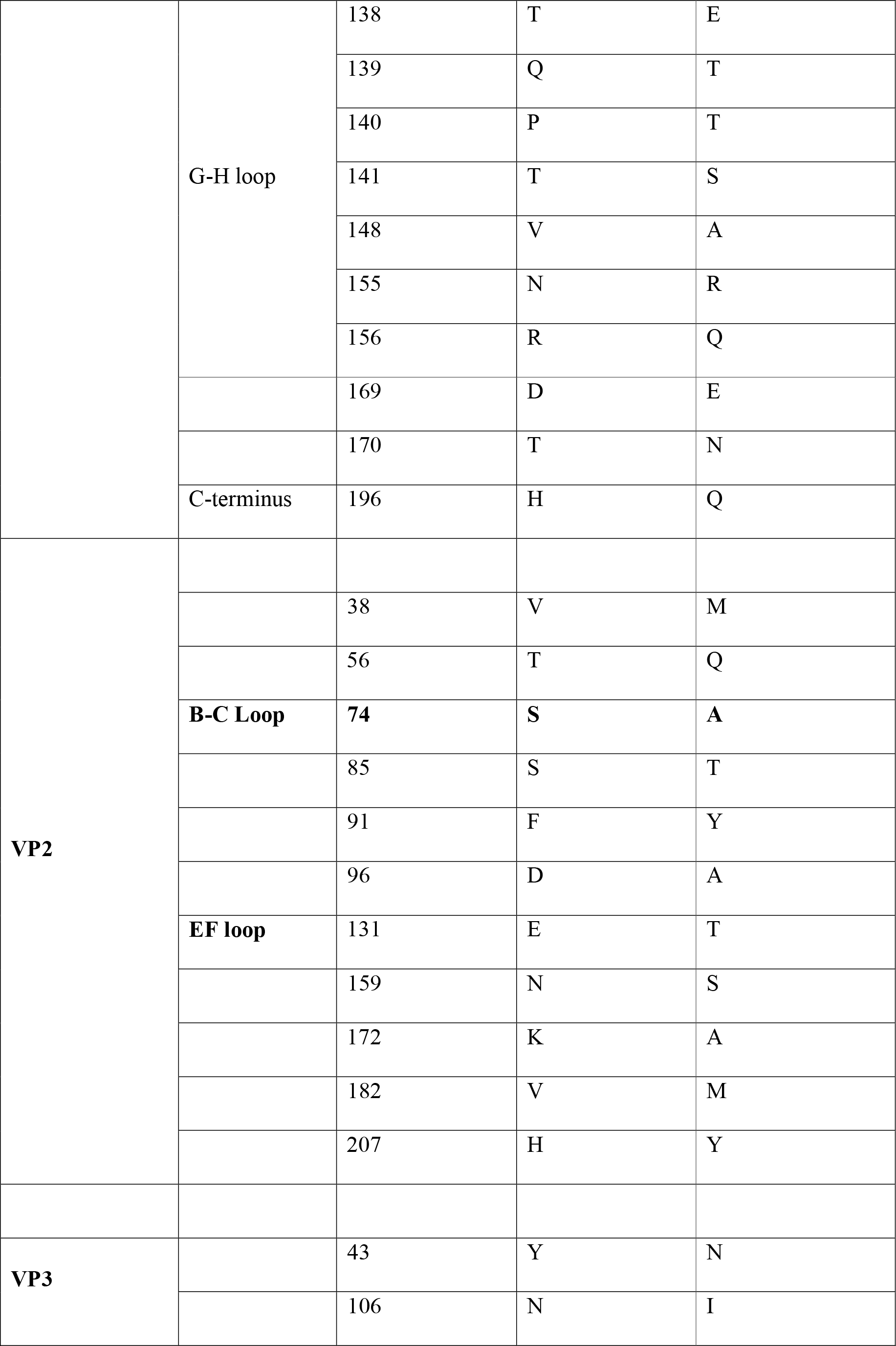
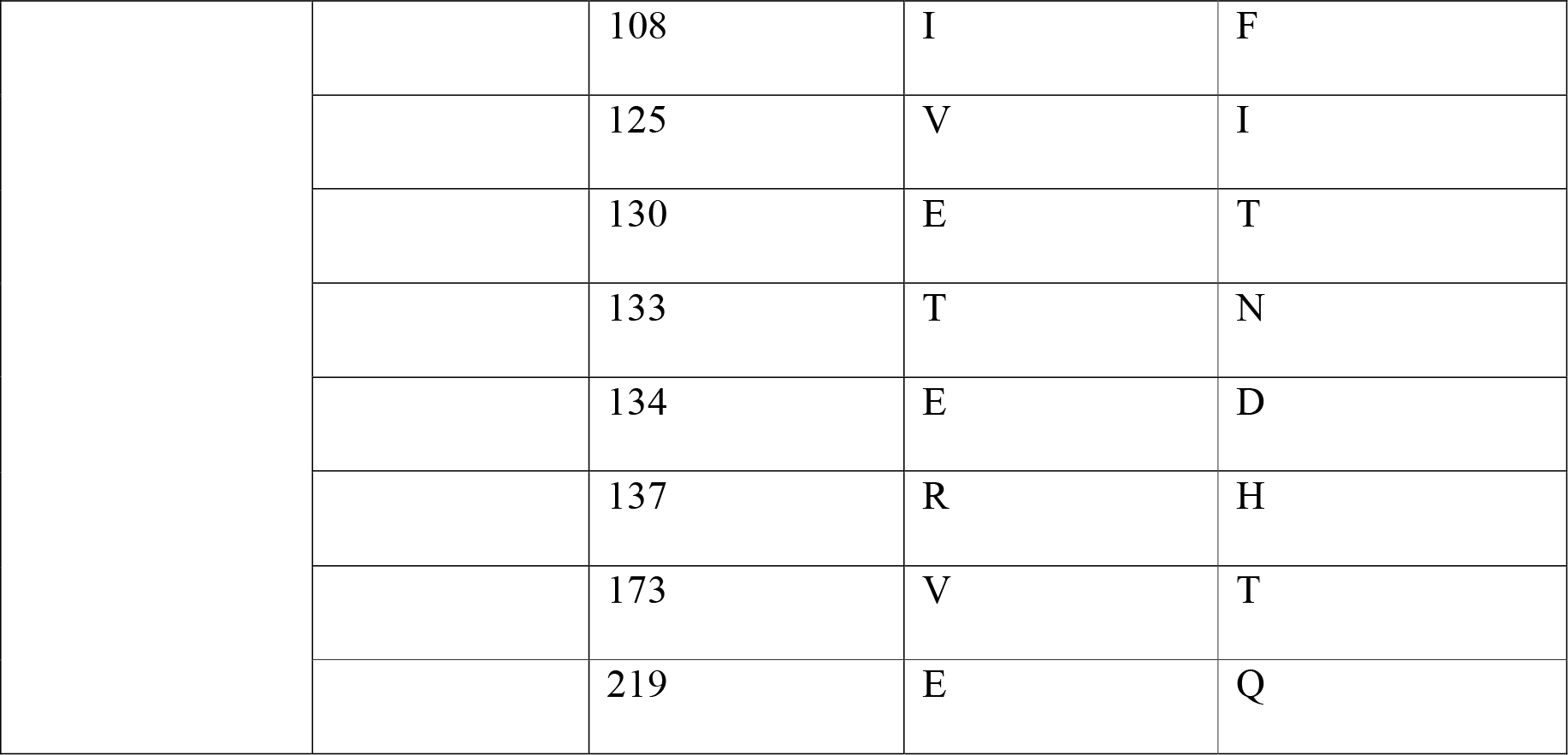
Amino acid mutation with respect to Reference vaccine strain.

**Table 2.** Percent divergence of in nucleotide sequence of the proteins between BAN/TA/Ma-167/2013 and BAN/DH/Sa-318/2018.

| Lpro | VP1 | VP2 | VP3 | VP4 | 2A | 2B | 2C | 3A | 3B | 3C | 3D |
| --- | --- | --- | --- | --- | --- | --- | --- | --- | --- | --- | --- |
| 87% | 91.3% | 91.3% | 92.3% | 90.2% | 87.5% | 94.2% | 93.2% | 93.7% | 93% | 93.3% | 94% |

The widespread genetic variations resulting in frequent emergence of the novel strain followed by rapid spreading, uncertainty in transmission, imprecise origin of FMD outbreaks, transboundary movement, possible intrusions into disease-free countries, poor surveillance scheme, and ineffective vaccination make the FMD control program complex in Bangladesh, which also pose a threat to disease-free countries. This complete genome characterization will help the scientific community to trace the outbreak source more precisely, check mutational pattern and possibly tackle the emergence of novel lineage by revealing the mystery hidden in the genome.

## Acknowledgements

The PI and co-authors thank the University Grants Commission (UGC), Government of the People’s Republic of Bangladesh for providing the fund for this work.

